# Early induction of cytokine release syndrome by rapidly generated CAR T cells in a preclinical mouse model

**DOI:** 10.1101/2022.11.12.515207

**Authors:** Naphang Ho, Arezoo Jamali, Angela Braun, Elham Adabi, Frederic B. Thalheimer, Christian J. Buchholz

## Abstract

Chimeric antigen receptor (CAR) T cells have emerged as effective strategy against B cell malignancies. Since the long manufacturing process limits patient accessability, short-term (st) CAR T cells are under investigation. Here, we evaluated CD19-CAR T cells 24 hours after exposure to lentiviral vectors. In co-culture with tumor cells and monocytes, stCAR T cells exhibited anti-tumoral activity and strong release of CRS-relevant cytokines (IL-6, IFN-γ, TNF-α, GM-CSF, IL-2, IL-10). When administered into tumor engrafted NSG-SGM3 mice, severe acute adverse events encompassing high body scoring, temperature and weight drop arised rapidly within 24 hours. Human (IFN-Y, TNF-α, IL-2, IL-10) and murine (MCP-1, IL-6, G-CSF) cytokines typical for severe cytokine release syndrome (CRS) were systemically elevated. Our data highlight potential safety risks of CAR T cells manufactured within short time and suggest simple models for their preclinical safety evaluation.

## Introduction

CAR T cells as new therapeutic modality with promising outcomes in hematologic malignancies have changed the cancer immunotherapy field vibrantly with rapid progress. The extensive efforts of the last two decades have resulted in outstanding achievments with several CAR T cell products having reached the market. However, a major drawback of the current CAR T cell products is the long manufacturing time, which can require weeks until the patient receives the therapeutic cell product.^1–3^

Therefore, the generation of CAR T cells within a few days is currently in focus of CAR T cell therapy research. The benefits of such CAR T cells would not only be an accelerated availability for patients, but potentially also a more advantageous cell composition of the CAR T cell product. For instance, CAR T cells generated within three days display a more naïve-like phenotype and exhibit superior proliferation and effector function post adoptive cell transfer into tumor engrafted mice in contrast to long-term expanded CAR T cells.^4^ While current studies reveal that short-term CAR T cells may provide a beneficial phenotype outperforming conventional CAR T cells by *in vivo* expansion and anti-tumor activity,^4–6^ the safety of these cells is less well characterized. Especially the short time period after exposure to vector particles prior to administration habors the likelihood that transduction is incomplete, making the properties of these short-term CAR T cells rather unpredictable. So far, preclinical studies focussed exclusively on the therapeutic activities of short-term CAR T cells.^4–7^ Assay systems identifying residual vector particles as well as a careful safety assessment of these CAR T cells are therefore urgently needed.

Cytokine release syndrome (CRS) and immune effector cell-associated neurotoxicity syndrome (ICANS) represent the most prominent side effects caused by conventional CAR T cells, which can reach fatal states in particular patients.^8^ Clinically, CRS is characterized by fever at the onset, followed by hypotension, capillary leak, and end-organ dysfunction with critical medical care requirements.^9^ The severity of CRS correleates to tumor burden, the amount of infused CAR T cells and their proliferation.^8, 10^ Commonly, CRS appears a few days after CAR T cell administration and peaks within a week. During this period high CAR T cell expansion can be observed, which comes along with massive elevation of various inflammatory cytokines in the serum, including IL-6, IL-10, IL-8, IFN-γ, and TNF-α.^8, 10^ Pre-clinical studies in mouse models showed that CRS is a complex interplay of activated CAR T cells encountering tumor antigens and further immune cells.^11, 12^ In particular, monocytes and macrophages have been identified as a major source for IL-6 and IL-1 and can therefore be regarded as key drivers for progression and induction of CRS.^11–15^ Although CRS can be managed by cytokine receptor blockers, e.g. tocilizumab, it remains in focus of ongoing research including the identification of predictive preclinical model systems.^11, 12, 14–16^

Here we focussed on the preclinical safety evaluation of short-term CAR T cells (stCAR T cells), which were generated within three days after T cell isolation. We show that a high percentage of stCAR T cells contain vector particles on their surface which disappear after prolonged cultivation. Their cytotoxic activity was confirmed in an *in vitro* killing assay, and in the presence of monocytes the secretion of CRS-relevant cytokines was observed. Upon administration into tumor engrafted NSG-SGM3 mice stCAR T cells rapidly induced CRS-related severe symptoms as well as release of high cytokine levels.

## Results

### Short-term CAR T cells are vector-bound cells with beneficial cell composition

For the generation of CD19-specific stCAR T cells human PBMCs were activated for two days with anti-CD3 and anti-CD28 activation antibodies in the presence of IL-7 and IL-15 and subsequently incubated with VSV-LV (MOI 4-5) for 24 hours. For characterization, stCAR T cells were analyzed by flow cytometry to detect the VSV glycoprotein (VSV-G) and myc-tag of the CAR. Most of the cells (about 50%) were single-positive for VSV-G while a small proportion of about 15% was additonally positive for the CAR (Figure 1A-B). Less than 5% of the cells were solely positive for the CAR with no detectable VSV-G (Figure 1A-B). To see if the vector-bound cells convert to CAR expressing T cells, the cells were further activated and cultivated. After three days, about 50% of the cells were single positive CAR T cells with no detectable VSV-G protein (Figure 1A-B). This demonstrates the incomplete but ongoing transduction in stCAR T cells.

**Figure 1:**
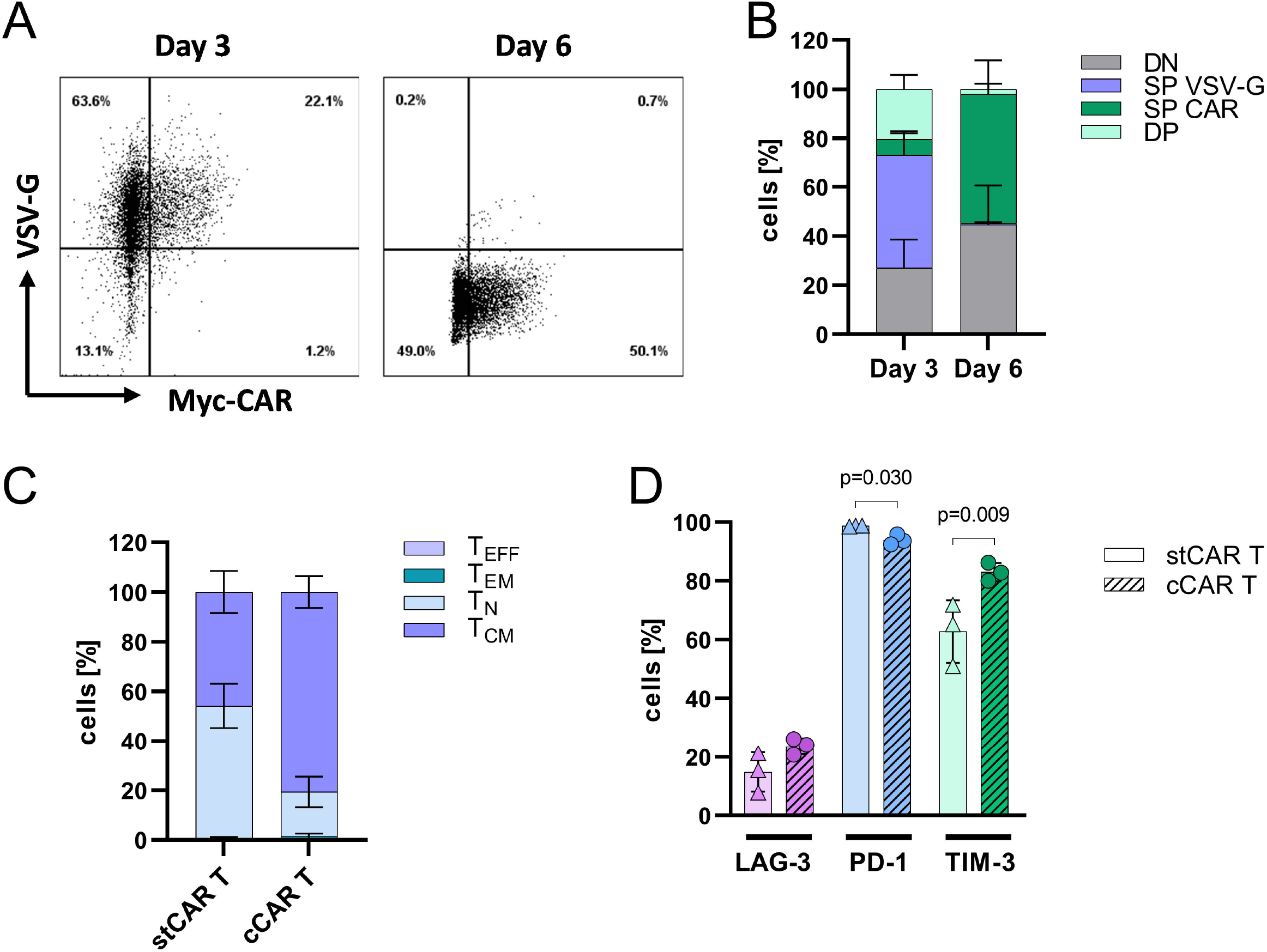
Characterization of short-term CAR T cells. Short-term CAR T cells (stCAR T) were characterized for presence of vector particle components and CAR expression as well as for phenotype and exhaustion markers using flow-cytometry. **A)** Representative dot plots for vector and CAR signal on stCAR T cells (Day 3) and after further activation and cultivation (Day 6) with indicated cell frequencies inside the quadrants. Gates were set according to the non-transduced negative control. **B)** Stacked bar diagrams summarize stCAR T cells on day 3 and day 6 with single positive VSV-G (SP VSV-G), single positive CAR (SP CAR), double positive (DP) and double negative (DN) cell frequencies. Phenotypes, determined by CD62L and CD45RA expression, **(C)** and exhaustion marker expression **(D)** of stCAR T cells were compared to conventional generated CAR T cells (cCAR T). Mean and standard deviation are presented for four donors in **(B)** and three donors in **(C, D)**. The statistic in **(D)** was determined by unpaired t-test with indicated significant p-values.

Further analysis revealed a beneficial cell composition of stCAR T cells compared to conventional CAR T cells (cCAR T cells), which were generated by three days of activation followed by three days of viral vector transduction. Both CAR T cells types were similar in CD3^+^ levels while viability was slightly reduced for stCAR T cells (Suppl. Fig. 1). Additionally, stCAR T cells showed a less differentiated phenotype with mainly naïve T cells (53%), whereas cCAR T cells were mostly represented by central memory T cells (80%) (Figure 1C). Moreover, exhaustion markers were generally less expressed on stCAR T cells with slightly lower LAG-3 and significantly reduced TIM-3 expression, although PD-1 expression was marginaly higher compared to cCAR T cells (Figure 1D).

This observation showed that stCAR T cells are rather vector particle-bound cells with a less differentiated and less exhausted phenotype.

### Short-term CAR T cells are cytotoxic active and induce CRS-relevant cytokines

To evaluate the cytotoxicity of stCAR T cells including their capability for CRS induction, a killing assay co-culture was designed to recapitulate key immune cell components involved in cytokine release. A co-culture of stCAR T cells and NALM6 tumor cells was supplemented with 10% monocytes. To avoid alloreactivity, monocytes and PBMCs, used for stCAR T cell generation, were derived from the same donor.

StCAR T cells displayed pronounced killing of tumor cells after one day of co-culture compared to control T cells with less than 20% remaining viable tumor cells (Figure 2A). At this time-point about 60% of T cells were CAR positive with a two-fold higher frequency for CD8 negative than CD8 positive T cells (Figure 2B). Thus, vector particle-bound T cells had converted to CAR T cells during the co-culture, which resulted in efficient tumor cell killing. Importantly, cytotoxicity and CAR expression were not influenced by the addition of monocytes to the culture (Figure 2A-B).

**Figure 2:**
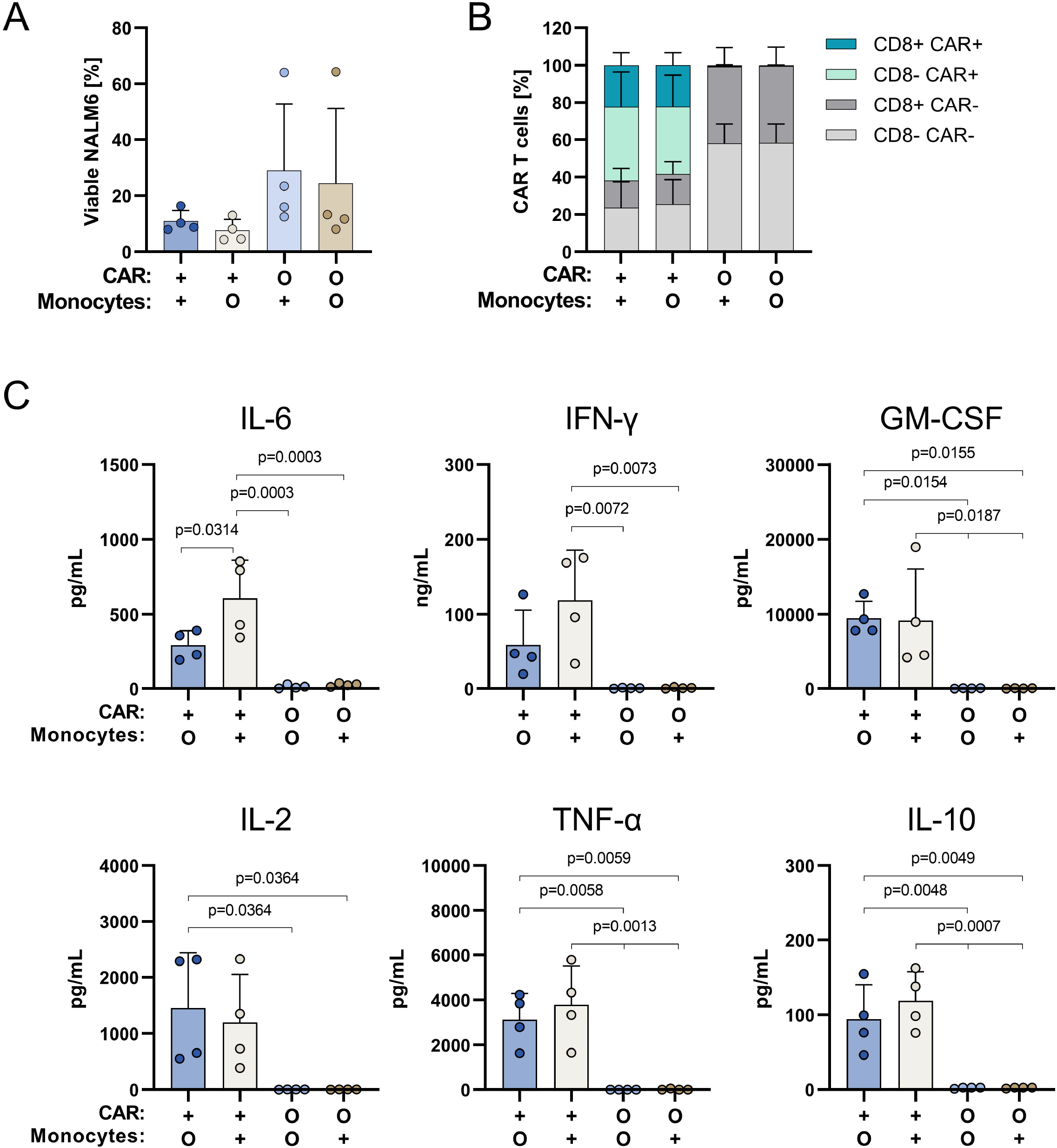
*In vitro* cytotoxicity assay and cytokine release. Short-term CAR T cells (+) or untransduced T cells (O) were co-cultured with CD19+ NALM6 cells at a 5:1 (effector: target) ratio in the presence (+) or absence (O) of 10% monocytes for 24 – 26h. Bar diagrams show cytotoxicity determined as the viability of tumor cells **(A)** and CAR expression on short-term CAR T cells **(B)** after co-culture by flow cytometry. **C)** Co-culture supernatant was analyzed for human cytokines with a bead-based multiplex kit. Data are shown for four donors with the mean and standard deviation of the group from two independent experiments. Statistics were determined by one-way ANOVA and Tukey’s multiple comparisons test with indicated significant p-values.

Analysis of human cytokines secreted during killing revealed tremendously elevated levels of CRS-associated pro-inflammatory IL-6, IFN-γ, GM-CSF, IL-2 and TNF-a in the cell culture supernatant compared to T cell control (Figure 2C). Strikingly, the clinical CRS-relevant cytokine IL-6 was significantly enhanced especially when monocytes had been supplemented to the co-culture (Figure 2C). Similarily, IFN-γ levels were increased by two-fold in the presence of monocytes and over 70-fold compared to the T cell control. Irrespectively of the addition of monocytes, GM-CSF, IL-2 and TNF-α were remarkably elevated in the supernatant. Notably, the anti-inflammatory IL-10, was also substantially elevated for stCAR T cells (Figure 2C). Overall, a high cytokine release was observed during anti-tumoral activity with induction of clinically relevant CRS-related cytokines in the presence of monocytes.

### CRS tumor model with short-term CAR T cells

Next, a CRS mouse model was set up to further evaluate stCAR T cells for their potential to induce systemic adverse events. Since the amount of target cells is related to the CAR T cell activity and hence the cytokine release, a high tumor load was chosen to achieve a sensitive CRS model. Accordingly, immunodeficient NSG-SGM3 mice were engrafted with luciferase positive tumor cells for 10 days before stCAR T cells were injected (Figure 3A). Tumor growth was carefully monitored by *in vivo* bioluminescence imaging system (IVIS), showing that tumor cells were located mainly in bones of the legs, hips and sternum (Suppl. Fig. 2A). One day before stCAR T cell injection, mice were distributed into the experimental groups based on equal tumor load (Suppl. Fig. 2B). Afterwards, 1×10^7^ stCAR T cells (determined as VSV-G^+^ and CAR^+^ cells) were administered intravenously. T cells activated and cultured under identical conditions were injected as control. Mice were then monitored tightly for overall health conditions, including appearance, animal activity, weight, temperature and plasma cytokine measurment.

**Figure 3:**
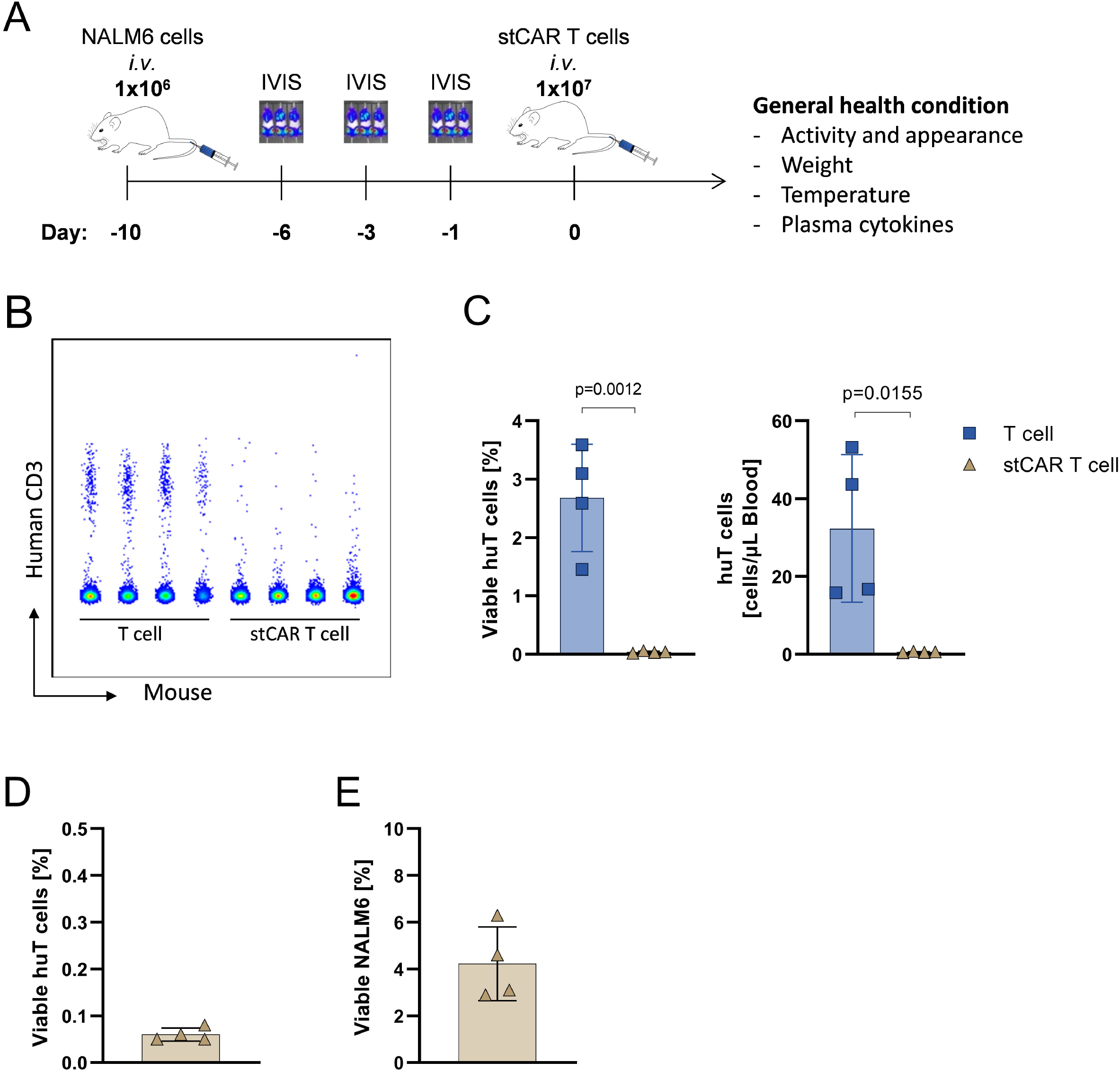
Evaluation of short-term CAR T cells in NSG-SGM3 mice. **A)** NSG-SGM3 mice were engrafted intraveneously (*i.v*.) with 1 ×10^6^ EBFP and luciferase expressing NALM6 tumor cells. Tumor load was monitored by *in vivo* bioluminescence imaging system (IVIS). 1 ×10^7^ short-term (st) CAR T cells or equal amount of T cells were injected *i.v*. on day 0 and mice were monitored for general health condition until termination criteria were reached. Human T cells in the blood one day after stCAR T cell or T cell administration were determined by flow-cytometry. **B)** Dot plot of human CD3+ T cells within all viable cells for each mouse. **C)** Bar diagrams show frequency of human T cells within viable cells and absolute quantification of human T cells per μL blood. Bone marrow cells were analyzed for frequency of human T cells **(D)** and tumor cells **(E)** in stCAR T cell treated mice on termination day. Single data point represents one mouse including mean and standard deviation of the group. n = 4 (T cell), 4 (stCAR T cell). Statistics were determined by unpaired t test with indicated significant p-values.

On the next morning, only mice treated with stCAR T cells showed strong ruffled fur, squinted eyes and reduced responsiveness. Interestingly, flow cytometry analysis of peripheral blood readily detected human T lymphocytes in the control group (3% of all viable cells corresponding to 35 cells per μL blood) but showed a clear absence of these cells in the blood of stCAR T cell-treated mice (Figure 3B-C). In bone marrow of stCAR T cell injected mice, 4% of EBFP expressing NALM6 as well as a few events of human T cells with a frequency below 0.05% of all viable cells could be detected (Figure 3D-E).

For the general health assessment, visual appearance, cage activity and weight loss were scored in a body index, which represents adverse effects with increasing scoring numbers. All mice treated with stCAR T cells exhibited high scoring one day after administration, whereas the T cell treated mice showed a similar scoring as naive mice (Figure 4A). Only after 12 days, the scoring index of T cell treated mice increased due to the tumor load (Figure 4A, Suppl. Figure 2C). To dissect the scoring in detail, relative weight loss and temperature changes to the start of the experiment were calculated. Correlating to the body scoring all mice showed a drop in weight on the day of termination (Figure 4B). Mice treated with stCAR T cells showed on average about 10% weight loss within 24h, whereas T cell treated mice experienced relative weight loss of about 5% only on day 12 (Figure 4B). Strikingly, only stCAR T cell treated mice displayed a very distinct temperature drop of over 2°C from the baseline within 24 hours (Figure 4C).

**Figure 4:**
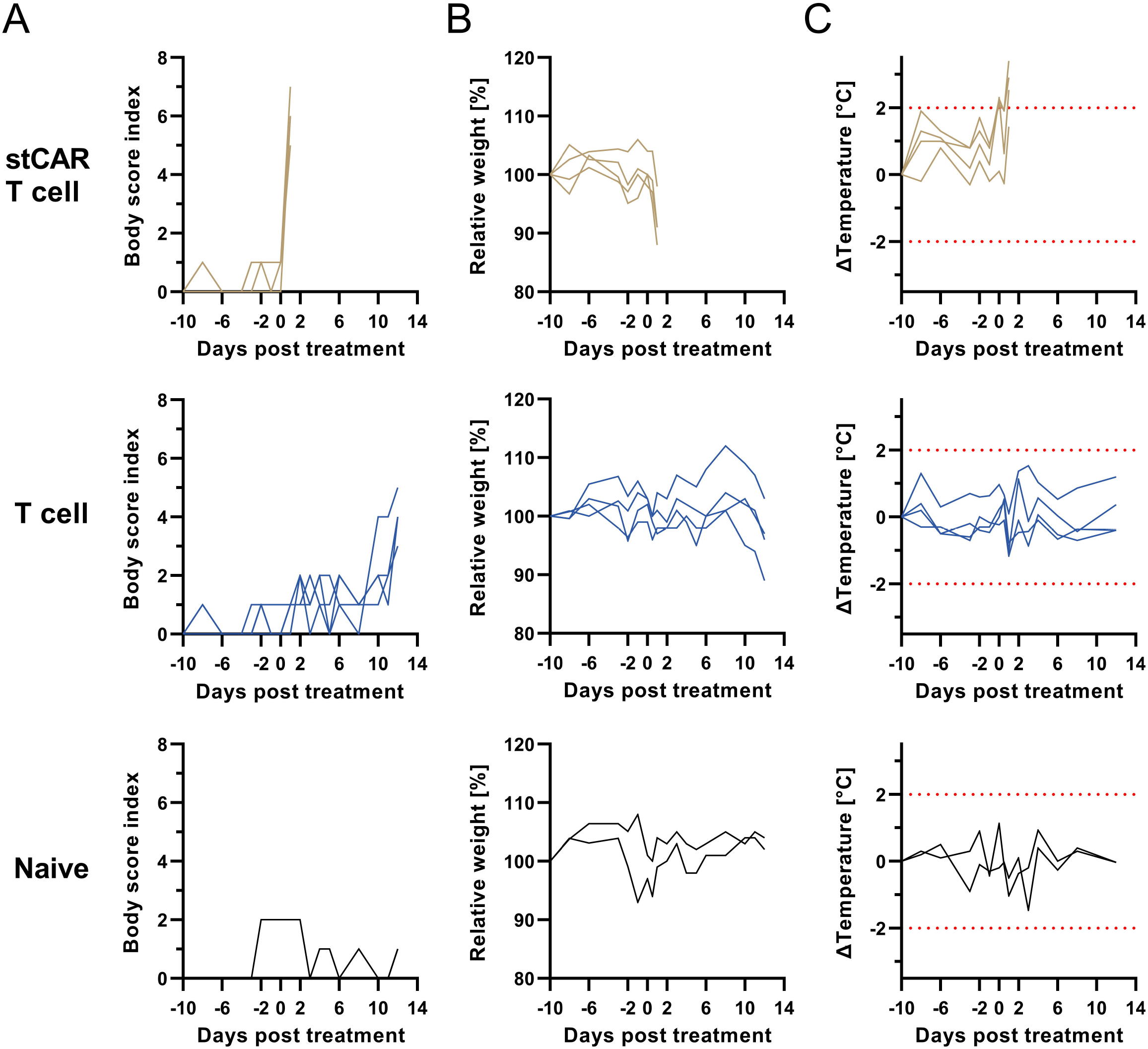
Acute adverse effects after short-term CAR T cell administration. NSG-SGM3 mice were engrafted intraveneously (*i.v*.) with 1 ×10^6^ EBFP and luciferase expressing NALM6 tumor cells. Afterwards, 1 ×10^7^ short-term (st) CAR T cells or identical T cell numbers were injected *i.v*. on day 0. Naïve mice did not receive tumor cells or T cells. Mice were monitored for general health condition, weight and temperature changes. **A)** Body scoring was assessed based on visual appearance, cage activity and weight loss of the mice.Weight was normalized to start weight at day −10 **(B)** and temperature change was determined as delta related to experiment start **(C)**. Red dotted lines indicate threshold of determined temperature change as adverse event. n= 2 (naive), 4 (T cell), 4 (stCAR T cell).

### Acute cytokine release after short-term CAR T cell administration

To verify if CRS was causative for the early mortality of stCAR T cell treated mice, typical CRS-associated human cytokines were assessed in the plasma on the day of termination. Blood of mice of the T cell group obtained on the same day provided xenogeneic baseline activities. Results showed a very high and significant increase in various CRS-related human cytokines in all mice treated with stCAR T cells compared to the T cell controls. In particular, IFN-γ was more than 130-fold increased in the stCAR T cell group reaching on average 70 ng/mL in plasma. Similarly, other CAR T cell associated pro-inflammatory cytokines, such as TNF-α (114-fold) and IL-2 (6-fold) and the anti-inflammatory cytokine IL-10 (54-fold) were substantially increased (Figure 5A). Interestingly, monocyte associated cytokines, such as IL-1β were not notably elevated, but on the other hand IL-6, a major CRS-relevant cytokine, was significantly increased, but at a lower level (<10 pg/mL) (Figure 5A).

**Figure 5:**
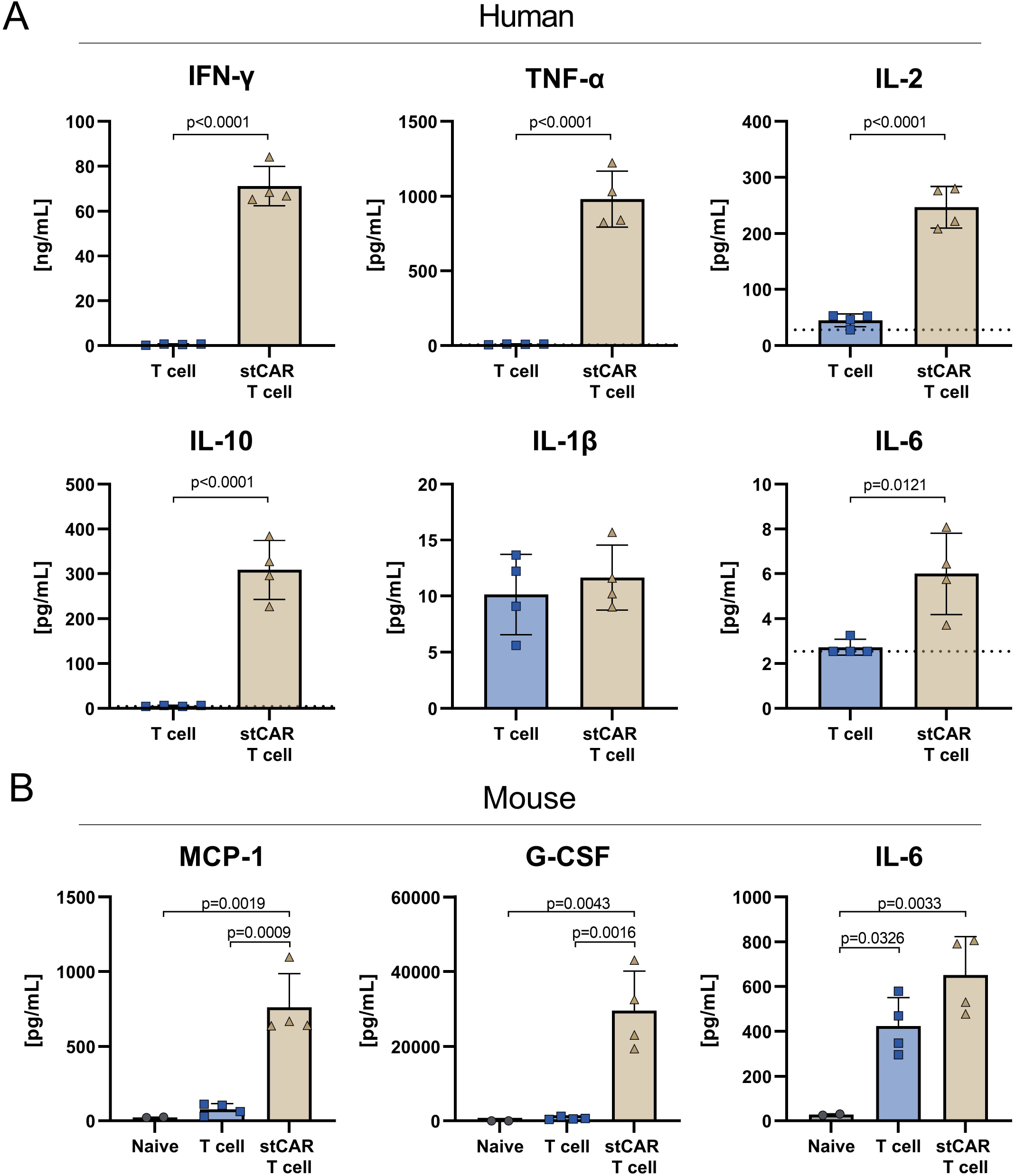
Cytokine release syndrome after short-term CAR T cell administration. **A)** Plasma was collected from both groups one day after treatment and human plasma cytokines were analyzed by bead-based multiplex human cytokine kit. **B)** Plasma was collected on the respective termination day of the groups, which was one day after treatment for the short-term (st) CAR T cell group and day 12 after treatment for the T cell and naive group and analyzed by bead-based multiplex murine cytokine kit. Bar digrams show cytokine for each mouse with mean and standard deviation of the group. Dotted line represents lower detection limit in the cytokine assay. n=2 (naive), 4 (T cell), 4 (stCAR T cell). Statistics were aquired by unpaired test in **(A)** and one-way ANOVA with Tukey’s multiple comparisons test in **(B)** with indicated significant p-values.

Due to the multicellular interplay of the whole immune system leading to systemic adverse events during CRS, the residing murine immune cells in the NSG-SGM3 model might have also been involved in the cytokine release syndrome process. In particular, this refers to murine myeloid cells including monocytes, macrophages, DCs and neutrophils all of which were detected in the spleen of NSG-SGM3 mice (Suppl. Figure S3). Interestingly, the majority of murine splenocytes were neutrophils in stCAR T cell treated mice, whereas T cell treated mice showed a more prominent presence of myeloid DCs on the final day (Suppl. Figure S3). To examine if these cells also contributed to the ongoing CRS, multiple murine cytokines were measured in the plasma of stCAR T cell and T cell treated mice on the respective termination days. Results show that especially murine MCP-1 and G-CSF were significantly elevated in stCAR T cell treated mice with about 9-fold and 59-fold increase compared to the other groups. Interestingly, murine IL-6 was elevated in both, the CAR T cell and T cell treated mice, compared to the naïve mice, but notably more in the CAR T cell group (Figure 5B). Particularly MCP-1 and G-CSF, but also the slightly enhanced release of IL-6 in stCAR T cell treated mice argue for systemic CRS induced acute morbidity.

### Discussion

Current approaches to refine CAR T cell therapy focus on the reduction of manufacturing times, down from weeks to few days or even hours. Due to the nature of CAR T cells being manufactured from patient-derived T lymphocytes, such stCAR T cells have to be considered as novel type of product potentially harboring specific safety risks. Accordingly, stCAR T cells require independent testing for activity and safety. Here, we established and evaluated assay systems for this task. The CD19-directed stCAR T cells resembled a less differentiated and exhausted phenotype when compared to conventional CAR T cells as previously described.^4–6^ Interestingly, our assay for the detection of vector components in these cells revealed that stCAR T cells consisted mainly of vector particle-bound cells with only a small fraction of cells being positive for the CAR. Only upon further cultivation and activation CAR becomes cell-surface expressed and vector particles dissapear, demonstrating that transduction is not fully completed in stCAR T cells. Previous studies did not track the presence of vector components, while CAR transferred as protein from packaging cells to transduced cells has been studied and detected,^7, 17, 18^ which however, did not mediate killing when reverse transcription and genome integration of the CAR gene was inhibited.^7^ Using an anti-VSV-G antibody our study allowed a more accurate characterization of stCAR T cells with respect to residual vector particles. A safety concern that comes along with administration of large amounts of LV-bound cells is gene delivery to off-target cells. Detachment of cell surface-bound vector particles from stCAR T cells or even binding of stCAR T cells to other cells, can result in undesired gene transfer mediated by VSV-G protein. If this occurs with patient tumor cells, fatal consequences may result.^19^ Accordingly, careful quality control of stCAR T cells must be ensured for a safe product.

A surprising and highly relevant outcome from our study was the fast induction of typical CRS symptoms within only 24 hours after stCAR T cell administration. This included strong ruffled fur, squinted eyes, reduced responsiveness, weight and temperature drop as well as high levels of plasma cytokines. The pattern of increased human plasma cytokines (IFN-γ, TNF-α, IL-2 and IL-10) was in line with CRS observed in patients and noteably also with the data from the *in vitro* cytotoxicity assay. Although, human monocytes were not present in the *in vivo* mouse model, CRS-induced adverse events by stCAR T cells could be sensitively captured. Besides human cytokines, elevated murine cytokines, including IL-6, G-CSF and MCP-1, underlined the ongoing multicellular interplay of human and murine cells in these mice during CRS. Immunophenotyping of residual murine immune cells in splenocytes of NSG-SGM3 mice revealed the presence of macrophages and monocytes, which presumably released murine IL-6. Further, murine MCP-1, a chemoattractant for monocytes, and murine G-CSF, an activator for granulocytes, were particularly increased in CRS affected mice. Both these cytokines are secreted by activated endothelial cells,^20, 21^ which were described for CD19-CAR T cell treated patients with severe CRS and ICANS.^22^ In addition, G-CSF is known to induce and immobilize neutrophil production and release from the bone marrow into the periphery^23, 24^ thus explaining the pronounced presence of murine neutrophils we observed in spleens of stCAR T cell-treated mice.

Such rapid kinetic for CRS induction has not been reported so far for conventionally manufactured CAR T cells in other pre-clinical CRS mouse models, where adverse events appeared between 2 – 5 days post CAR T cell administration.^11, 12, 16^ Interestingly, similar rapid kinetics have been observed in patients with life-threatening grade of CRS (grade 4 or more) within 25 hours post conventional CAR T cell infusion.^25–27^ In line with the rapid onset of CRS, stCAR T cells, in contrast to control T cells, had almost completely vanished from blood after one day. Detection of human T cells in the bone marrow of these mice suggests rapid migration of stCAR T cells to the tumor-residing niche which may have promoted the fast and severe CRS. This feature has recently been attributed especially to rapidly manufactured CAR T cells and was correlated to enhanced expression of the chemokine receptor CXCR4.^6^

Current preclinical data about CAR T cells with shortened manufacturing time tested in therapeutical mouse models did not reveal any concerncs regarding CRS and adverse events so far.^4–7^ While CAR T cell generation followed a similar protocol, with respect to the short lentiviral vector exposure to the cells, relevant differences to our study refer to the mouse strain (NSG and NOG) and the administered CAR T cell doses, which were 3 – 10 fold lower. Indeed, the NSG-SGM3 mouse model, which was only used in our study, has previously proven to be more sensitive for CRS than the parental NSG strain when assessing the safety of anti-CD28 superagonist and anti-PD1 monoclonal antibodies.^28^ While we cannot exclude that besides the mouse strain the higher administered dose contributed to rapid CRS induction, so far the only data from clinical trials with rapidly manufactured CD19-CAR T cells documented CRS occurrence about one week post treatment.^5, 6^ Note, that this clinical study administered a rather low CAR T cell dose (10^4^ to 10^5^ cells/kg body weight), yet induced severe CRS in some patients. In comparison, we administered approximately 5×10^8^ cells/kg and thus a more than 1000-fold higher dose, which was well in the range of what was applied in CRS mouse models for conventional CAR T cells.^11, 12, 16^ Nevertheless, our results call for careful safety assessement of stCAR T cells and suggest that dose administration in the range of 10^7^ cells/kg as used with conventional CAR T cells^25, 29, 30^ may not be safe for rapidly manufactured CAR T cells. Further preclinical assessment using the CRS model presented in this study will give a more comprehensive understanding about the dosage regimen and anti-tumoral activity for this novel type of CAR T cell product.

## Material and Methods

### Vector production

The protocol for production and characterization of VSV-LV stocks was adapted from previously.^31^ VSV-LV contained the transfer plasmid pS-CD19-CAR-W encoding a second generation CD19-CAR with a CD28 and CD3z signaling domain under control of the SFFV promotor. In brief, 2.5 × 10^7^ β2M^-/-^, CD47^high^ HEK293T cells were seeded in a T175 flask 24 h before transfection with 17.5 μg of packaging plasmid pCMVdR8.9, 11.4 μg of transfer plasmid pSEW-mycCD19.CAR(28z) and 6.1 μg of envelope plasmid pMDG-2. After 2 days the vector particles released into the supernatant were collected, concentrated, and purified through a 20% sucrose cushion (4500 × g, 24 h at 4°C). The supernatant was discarded, the pellet was resuspended in 60 μL of PBS, and the vector stocks were stored at −80°C. Gene transfer activity of the vector stocks was determined by transduction of 8 × 10^4^ activated human PBMCs in a serial dilution of the vector stocks. PBMCs were transduced via spinfection (at 850 × g, 90 min and 32°C). Transducing units were calculated based on the linear range of the gene transfer activity after 3 days. To determine particle numbers, nanoparticle tracking analysis was performed at the NanoSight NS300 (Malvern Panalytical).

### Primary cells and cell lines

Human PBMCs were isolated from freshly sampled human blood, purchased from the German blood donation center (DRK-Blutspendedienst, Frankfurt am Main) by Histopaque gradient centrifugation. Primary cells were cultured in 4Cell^®^ Nutri-T medium (Sartorius) supplemented with 0.4% penicillin/streptomycin, 25 U/mL human IL-7 (Miltenyi Biotec) and 50 U/mL human IL-15 (Miltenyi Biotec). Monocytes were isolated from PBMCs by anti-human CD14 (REA599, APC, Miltenyi Biotec) antibody labeling and subsequent isolation using a magnetic anti-APC-Microbead Kit (Miltenyi Biotec). NALM6 cells were cultivated in RPMI 1640 medium (Biowest) supplemented with 10% fetal bovine serum (FBS) (Biochrom AG) and 2 mM glutamine (Sigma-Aldrich). β2M^-/-^, CD47high HEK293T cells were cultivated in DMEM (Gibco) supplemented with 10% FBS and 2 mM glutamine. All cells were cultivated at 37°C, 5% CO2, and 90% humidity.

### Conventional CAR T cell generation

PBMCs were activated in plates coated with 1 μg/mL anti-CD3 mAb (clone: OKT3, Miltenyi Biotec) and cultured in the presence of 3 μg/mL anti-CD28 mAb (clone: 15E8, Miltenyi Biotec), 25 U/mL human IL-7 (Miltenyi Biotec) and 50 U/mL human IL-15 (Miltenyi Biotec) for three days. Afterward, 8 × 10^4^ activated human PBMCs were transduced with 0.5 μL VSV-LV (MOI 4-5) via spinfection (at 850 × g, 90 min and 32°C) and further cultivated for three days.

### Short-term CAR T cell generation

PBMCs were activated in plates coated with 1 μg/mL anti-CD3 mAb and cultured in the presence of 3 μg/mL anti-CD28 mAb, 25 U/mL human IL-7 and 50 U/mL human IL-15 for two days. Next, 8 × 10^4^ activated human PBMCs were incubated with 0.5 μL VSV-LV (MOI 4-5) in a flat-bottom 96-well plate or respectively upscaled for bigger production in a 24-well plate. Thereafter cells were cultivated for 24 hours until short-term CAR T cells were harvested, washed and used for subsequent experiments. To evaluate transition of short-term CAR T cells to fully CAR expressing T cells, short-term CAR T cells were seeded in 1 μg/mL anti-CD3 mAb precoated plates and cultured in the presence of 3 μg/mL anti-CD28 mAb, 25 U/mL human IL-7 and 50 U/mL human IL-15 for three further days.

### Monocyte supplemented cytotoxicity assay

5 × 10^4^ short-term CAR T cells (determined as VSV-G^+^ and CAR^+^ cells) were co-cultured with 1×10^4^ NALM6 cells prelabeled with CellTrace^®^ Violet (CTV) (ThermoFisher) in 200 μL 4Cell^®^ Nutri-T medium (Sartorius) supplemented with 0.4% penicillin/streptomycin in a flat-bottom 96-well plate. In addition, co-culture was further supplemented with 5 × 10^3^ isolated autologous monocytes or without monocytes. After 24 – 26h supernatant was taken, centrifuged at 300 × g for 5 min, and transferred into a new 96-well plate and stored at −20°C until cytokine measurement. Cytotoxicity was determined as the percentage of viable cells within CTV positive cells. CAR T cell level was assessed within the CTV negative and CD3+ population by flow cytometry.

### Cytokine assay

Cytokines in frozen cell culture supernatant or frozen plasma were analyzed with a customized multiplex human or mouse cytokine Legendplex kit (BioLegend). Samples were measured at the MACS Quant Analyzer10 (Miltenyi Biotec) and analyzed with the Legendplex software (v.8.0, BioLegend).

### CRS tumor mouse model

All animal experiments were performed in accordance with the regulations of the German animal protection law and the respective European Union guidelines.

NSG-SGM3 (NOD.Cg-Prkdcscid Il2rgtm1Wjl Tg(CMV-IL3,CSF2,KITLG)1Eav/MloySzJ, ID: 013,062) mice were purchased from Jackson Laboratory. To establish a CRS tumor model, mice were engrafted for 10 days with 1 × 10^6^ EBFP and firefly luciferase expressing NALM6 tumor cells by intravenous (*i.v*.) injection. The tumor burden of the mice was monitored twice a week using *in vivo* imaging (IVIS Spectrum, Perkin Elmer). For this purpose, 150 μg/g of body weight D-luciferin (Perkin Elmer) was injected intraperitoneally and imaging data was obtained 10 minutes later. One day prior to short-term CAR T cell administration, mice were imaged and animals were arranged into two groups for equal distribution of tumor engraftment. One day later 1 × 10^7^ short-term CAR T cells, defined as vector-particle and CAR positive cells, were injected *i.v*. As a control, the respective number of T cells activated and cultivated under the same condition were injected. Naïve mice, which did not receive tumor cells or T cells, were included to assess background signals in IVIS imaging and as a negative control for murine cytokine analysis. Animals were monitored for general health parameters, including visual appearance, cage activity and weight change. In addition, the temperature was measured in technical triplicates with an infrared thermometer at the anogenital area. Temperature change is presented as delta temperature, subtracted from the value prior experiment start on day −10. Changes of more than 2°C from the initial temperature value were defined as adverse events. Blood was taken one day after short-term CAR T cell administration. On the final day, blood and bone marrow were harvested for further analysis.

### Preparation of single cell suspension from organs

Blood and bone marrow were collected from each mouse. Plasma was separated from blood cells by centrifugation at 300 × g for 5 min and second centrifugation of the supernatant at 16000 × g for 5 min to remove residual cell debris. Plasma was then stored at −80°C until cytokine analysis. Bone marrow cells were harvested by opening ends of the bones and centrifugation at 4,600 × g for 3 min in perforated 0.5 mL tubes inside a fresh 1.5 mL tube containing RPMI medium. Next, the cell suspension was passed through a 70 μm cell strainer. Isolated single-cell suspension of the organ and the blood were washed with PBS and subsequently incubated in BD Pharm Lyse buffer (BD Biosciences) for 10 min for erythrocyte lysis. Isolated single-cell suspensions of the organs were used for flow cytometry analysis.

### Flow cytometry analysis

Staining for flow cytometry analysis was performed in PBS supplemented with 2% FBS. Primary cells were blocked with human FcR blocking reagent (Miltenyi Biotec) and mouse-derived samples were additionally blocked with mouse FcR-blocking reagent (Miltenyi Biotec). The following anti-human antibodies were used for FACS staining: CD45 (2D1, BV510, BioLegend), CD3 (BW264/56, PerCP, Miltenyi Biotec; HIT3a, BV605, BD Bioscience), Myc for CAR detection (SH1-26e7.1.3, FITC, Miltenyi Biotec), CD14 (REA599, APC, Miltenyi Biotec), CD4 (VIT-4, PerCp, Miltenyi Biotec, VIT-4, VioBlue, Miltenyi Biotec), CD8 (RPA-T8, BV786, BD Bioscience; BW135/80, APC, Miltenyi Biotec). To determine cell viability, fixable viability dye (eFluor 780, Thermo Fisher Scientific) was used. Stained samples were fixed in 1% PFA and stored at 4°C until measurement. For vector particle staining against the glycoprotein of VSV, samples were incubated with the primary mouse-derived anti-VSV-G antibody (8G5F11, unlabeled, Kerafest) for 15 min at 4°C and washed thoroughly twice. Afterwards, cells were stained with secondary anti-mouse IgG antibody (polyclonal, AF647, Jackson ImmunoResearch) for additional 15 min at 4°C and subsequently washed three times. Afterwards, normal antibody staining as described above was performed. Measurement was carried out by the LSR Fortessa (BD Biosciences) or MACS Quant Analyzer 10 (Miltenyi Biotec) and data were analyzed using FlowJo v.10.1 (BD Biosciences) or FCS Express 6 (De Novo Software). To determine absolute counts in the blood, defined volumes of 4 μm CountBright Plus absolute counting beads (Invitrogen) were added to the samples before flow cytometry measurement. Phenotype of CAR T cells was determined by CD45RA and CD62L expression. Naïve-like T cells were defined as CD45RA and CD62L positive, central memory T cells as CD62L positive, effector memory T cells as double negative and effector T cells as CD45RA positive. In mouse samples, tumor cells were gated on human CD45 negative cells and determined as EBFP positive cells. Human T cells were identified as human CD3 positive and EBFP negative cells.

### Body scoring

To assess and score general health condition of the animals, relative weight loss, cage activity and visual appearance were each scored separately from 0 to 3 and added together for the total body score. Healthy mice with no weight loss, normal cage activity and normal appearance were scored with 0. Animals with increasing weight loss (>0%; >5%; >10%), decreasing cage activity (less active; moderate active; barely responsive) and unhealthy visual appearance (slightly ruffled fur; ruffled fur with hunched back position; ruffled fur with hunched back position and squinted eyes) were scored accordingly (1; 2; 3). Mice were terminated when total body scoring was higher than 4 or with scoring 4 for more than two consecutive days.

### Statistical analysis

Data was analyzed using the GraphPad Prism software version 8 (Graph-Pad Software, USA). Statistical differences were assessed as indicated in the figure legend by using unpaired t-test, one-way and two-way ANOVA test with Tukey’s or Sidak’s multiple comparisons test. Differences were considered significant at p < 0.05.

## Data availability

All authors declare that data are available within the article or the supplemental information files.

## Acknowledgment

This work was supported by grants from the Bundesminsterium für Gesundheit (ZMV I 1 - 25 18 FSB 404), the Deutsche Krebshilfe (70114099) and the European Union’s Horizon 2020 research and innovation programme STACCATO under the Marie Skłodowska-Curie grant agreement (No. 813453.) to CJB.

## Author contributions

N.H. and F.B.T. designed the experiments. N.H., F.B.T. and A.J. performed the experiments and evaluated data. A.B. contributed to perform and evaluate data of the cytotoxicity assay and E.A. designed the protocol for vector particle detection. C.J.B and F.B.T. conceived and designed the study, acquired grants and supervised the work. N.H., A.J. and C.J.B wrote the manuscript.

## References

1. Levine, B.L., Miskin, J., Wonnacott, K., and Keir, C. (2017). Global Manufacturing of CAR T Cell Therapy. Molecular therapy. Methods & clinical development 4, 92–101.

2. Vormittag, P., Gunn, R., Ghorashian, S., and Veraitch, F.S. (2018). A guide to manufacturing CAR T cell therapies. Current opinion in biotechnology 53, 164–181.

3. Abou-El-Enein, M., Elsallab, M., Feldman, S.A., Fesnak, A.D., Heslop, H.E., Marks, P., Till, B.G., Bauer, G., and Savoldo, B. (2021). Scalable Manufacturing of CAR T cells for Cancer Immunotherapy. Blood cancer discovery 2, 408–422.

4. Ghassemi, S., Nunez-Cruz, S., O’Connor, R.S., Fraietta, J.A., Patel, P.R., Scholler, J., Barrett, D.M., Lundh, S.M., Davis, M.M., and Bedoya, F., et al. (2018). Reducing Ex Vivo Culture Improves the Antileukemic Activity of Chimeric Antigen Receptor (CAR) T Cells. Cancer immunology research 6, 1100–1109.

5. Zhang, C., He, J., Liu, L., Wang, J., Wang, S., Liu, L., Ge, J., Gao, L., Gao, L., and Kong, P., et al. (2022). Novel CD19 chimeric antigen receptor T cells manufactured next-day for acute lymphoblastic leukemia. Blood cancer journal 12, 96.

6. Yang, J., He, J., Zhang, X., Li, J., Wang, Z., Zhang, Y., Qiu, L., Wu, Q., Sun, Z., and Ye, X., et al. (2022). Next-day manufacture of a novel anti-CD19 CAR-T therapy for B-cell acute lymphoblastic leukemia: first-in-human clinical study. Blood cancer journal 12, 104.

7. Ghassemi, S., Durgin, J.S., Nunez-Cruz, S., Patel, J., Leferovich, J., Pinzone, M., Shen, F., Cummins, K.D., Plesa, G., and Cantu, V.A., et al. (2022). Rapid manufacturing of non-activated potent CAR T cells. Nature biomedical engineering 6, 118–128.

8. Morris, E.C., Neelapu, S.S., Giavridis, T., and Sadelain, M. (2022). Cytokine release syndrome and associated neurotoxicity in cancer immunotherapy. Nature reviews. Immunology 22, 85–96.

9. Lee, D.W., Santomasso, B.D., Locke, F.L., Ghobadi, A., Turtle, C.J., Brudno, J.N., Maus, M.V., Park, J.H., Mead, E., and Pavletic, S., et al. (2019). ASTCT Consensus Grading for Cytokine Release Syndrome and Neurologic Toxicity Associated with Immune Effector Cells. Biology of blood and marrow transplantation: journal of the American Society for Blood and Marrow Transplantation 25, 625–638.

10. Donnadieu, E., Luu, M., Alb, M., Anliker, B., Arcangeli, S., Bonini, C., Angelis, B. de, Choudhary, R., Espie, D., and Galy, A., et al. (2022). Time to evolve: predicting engineered T cell-associated toxicity with next-generation models. Journal for immunotherapy of cancer 10.

11. Giavridis, T., van der Stegen, S.J.C., Eyquem, J., Hamieh, M., Piersigilli, A., and Sadelain, M. (2018). CAR T cell-induced cytokine release syndrome is mediated by macrophages and abated by IL-1 blockade. Nature medicine 24, 731–738.

12. Norelli, M., Camisa, B., Barbiera, G., Falcone, L., Purevdorj, A., Genua, M., Sanvito, F., Ponzoni, M., Doglioni, C., and Cristofori, P., et al. (2018). Monocyte-derived IL-1 and IL-6 are differentially required for cytokine-release syndrome and neurotoxicity due to CAR T cells. Nature medicine 24, 739–748.

13. Barrett, D.M., Singh, N., Hofmann, T.J., Gershenson, Z., and Grupp, S.A. (2016). Interleukin 6 Is Not Made By Chimeric Antigen Receptor T Cells and Does Not Impact Their Function. Blood 128, 654.

14. Singh, N., Hofmann, T.J., Gershenson, Z., Levine, B.L., Grupp, S.A., Teachey, D.T., and Barrett, D.M. (2017). Monocyte lineage-derived IL-6 does not affect chimeric antigen receptor T-cell function. Cytotherapy 19, 867–880.

15. Sachdeva, M., Duchateau, P., Depil, S., Poirot, L., and Valton, J. (2019). Granulocyte-macrophage colony-stimulating factor inactivation in CAR T-cells prevents monocyte-dependent release of key cytokine release syndrome mediators. The Journal of biological chemistry 294, 5430–5437.

16. Arcangeli, S., Bove, C., Mezzanotte, C., Camisa, B., Falcone, L., Manfredi, F., Bezzecchi, E., El Khoury, R., Norata, R., and Sanvito, F., et al. (2022). CAR T cell manufacturing from naive/stem memory T lymphocytes enhances antitumor responses while curtailing cytokine release syndrome. The Journal of clinical investigation 132.

17. Cordes, N., Kolbe, C., Lock, D., Holzer, T., Althoff, D., Schäfer, D., Blaeschke, F., Kotter, B., Karitzky, S., and Rossig, C., et al. (2021). Anti-CD19 CARs displayed at the surface of lentiviral vector particles promote transduction of target-expressing cells. Molecular therapy. Methods & clinical development 21, 42–53.

18. Jamali, A., Kapitza, L., Schaser, T., Johnston, I.C.D., Buchholz, C.J., and Hartmann, J. (2019). Highly Efficient and Selective CAR-Gene Transfer Using CD4-and CD8-Targeted Lentiviral Vectors. Molecular therapy. Methods & clinical development 13, 371–379.

19. Ruella, M., Xu, J., Barrett, D.M., Fraietta, J.A., Reich, T.J., Ambrose, D.E., Klichinsky, M., Shestova, O., Patel, P.R., and Kulikovskaya, I., et al. (2018). Induction of resistance to chimeric antigen receptor T cell therapy by transduction of a single leukemic B cell. Nature medicine 24, 1499–1503.

20. Demetri, G.D., and Griffin, J.D. (1991). Granulocyte colony-stimulating factor and its receptor. Blood 78, 2791–2808.

21. Deshmane, S.L., Kremlev, S., Amini, S., and Sawaya, B.E. (2009). Monocyte chemoattractant protein-1 (MCP-1): an overview. Journal of interferon & cytokine research: the official journal of the International Society for Interferon and Cytokine Research 29, 313–326.

22. Gust, J., Hay, K.A., Hanafi, L.-A., Li, D., Myerson, D., Gonzalez-Cuyar, L.F., Yeung, C., Liles, W.C., Wurfel, M., and Lopez, J.A., et al. (2017). Endothelial Activation and Blood-Brain Barrier Disruption in Neurotoxicity after Adoptive Immunotherapy with CD19 CAR-T Cells. Cancer discovery 7, 1404–1419.

23. Ulich, T.R., Del Castillo, J., and Souza, L. (1988). Kinetics and mechanisms of recombinant human granulocyte-colony stimulating factor-induced neutrophilia. The American Journal of Pathology 133, 630–638.

24. Semerad, C.L., Liu, F., Gregory, A.D., Stumpf, K., and Link, D.C. (2002). G-CSF is an essential regulator of neutrophil trafficking from the bone marrow to the blood. Immunity 17, 413–423.

25. Maude, S.L., Frey, N., Shaw, P.A., Aplenc, R., Barrett, D.M., Bunin, N.J., Chew, A., Gonzalez, V.E., Zheng, Z., and Lacey, S.F., et al. (2014). Chimeric antigen receptor T cells for sustained remissions in leukemia. The New England journal of medicine 371, 1507–1517.

26. Teachey, D.T., Lacey, S.F., Shaw, P.A., Melenhorst, J.J., Maude, S.L., Frey, N., Pequignot, E., Gonzalez, V.E., Chen, F., and Finklestein, J., et al. (2016). Identification of Predictive Biomarkers for Cytokine Release Syndrome after Chimeric Antigen Receptor T-cell Therapy for Acute Lymphoblastic Leukemia. Cancer discovery 6, 664–679.

27. Hay, K.A., Hanafi, L.-A., Li, D., Gust, J., Liles, W.C., Wurfel, M.M., López, J.A., Chen, J., Chung, D., and Harju-Baker, S., et al. (2017). Kinetics and biomarkers of severe cytokine release syndrome after CD19 chimeric antigen receptor-modified T-cell therapy. Blood 130, 2295–2306.

28. Ye, C., Yang, H., Cheng, M., Shultz, L.D., Greiner, D.L., Brehm, M.A., and Keck, J.G. (2020). A rapid, sensitive, and reproducible in vivo PBMC humanized murine model for determining therapeutic-related cytokine release syndrome. FASEB journal: official publication of the Federation of American Societies for Experimental Biology 34, 12963–12975.

29. Turtle, C.J., Hanafi, L.-A., Berger, C., Gooley, T.A., Cherian, S., Hudecek, M., Sommermeyer, D., Melville, K., Pender, B., and Budiarto, T.M., et al. (2016). CD19 CAR-T cells of defined CD4+:CD8+ composition in adult B cell ALL patients. The Journal of clinical investigation 126, 2123–2138.

30. Neelapu, S.S., Locke, F.L., Bartlett, N.L., Lekakis, L.J., Miklos, D.B., Jacobson, C.A., Braunschweig, I., Oluwole, O.O., Siddiqi, T., and Lin, Y., et al. (2017). Axicabtagene Ciloleucel CAR T-Cell Therapy in Refractory Large B-Cell Lymphoma. The New England journal of medicine 377, 2531–2544.

31. Weidner, T., Agarwal, S., Perian, S., Fusil, F., Braun, G., Hartmann, J., Verhoeyen, E., and Buchholz, C.J. (2021). Genetic in vivo engineering of human T lymphocytes in mouse models. Nature protocols 16, 3210–3240.

