## Supplemental information for "Early induction of cytokine release syndrome by rapidly generated CAR T cells in a preclinical mouse model"

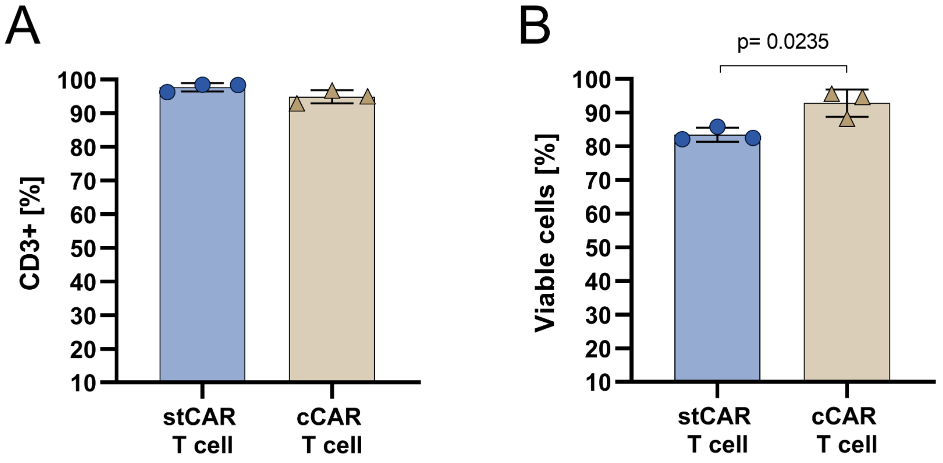


**Figure S1: CD3 expression and viability of short-term CAR T cells and conventional CAR T cells.**
Short-term (st) CAR T cells and conventional (c) CAR T cells were generated from human PBMCs. T cell frequency **(A)** and cell viability **(B)** after CAR T cell generation were determined by flow-cytometry. Mean and standard deviation are shown for three donors. Statistics were determined by unpaired t-test with indicated significant p-values.

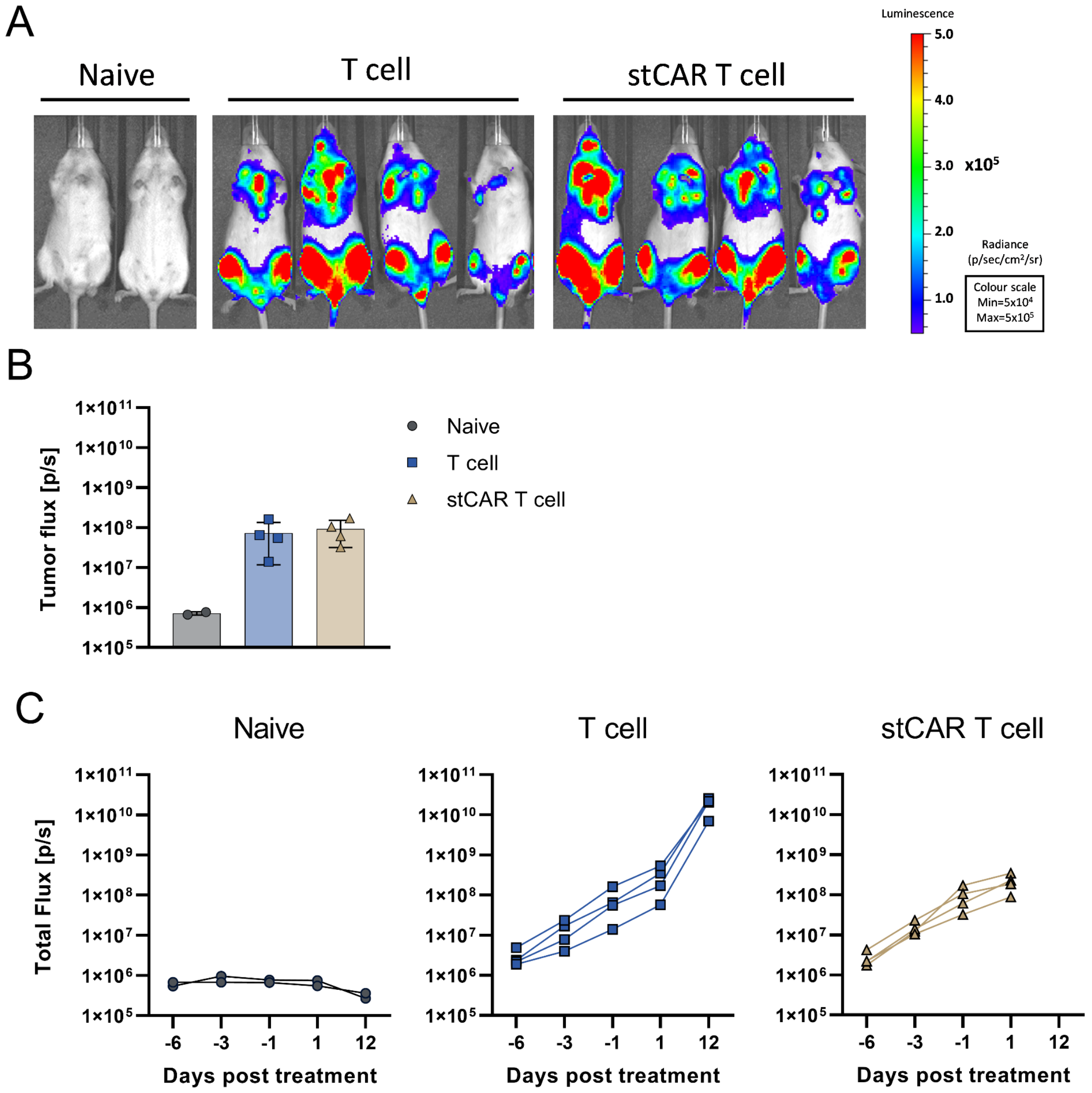


**Figure S2: Tumor cell engraftment in NSG-SGM3 mice.**

Tumor burden in NSG-SGM3 mice engrafted with EBFP and luciferase expressing NALM6 tumor cells were determined by *in vivo* bioluminescence imaging system (IVIS). Mice without tumor cell engraftment (naïve) were used to determine background signals. Tumor burden one day prior short-term (st) CAR T cell and T cell treatment are shown as IVIS images of the ventral side of each mouse with color scale for tumor signal intensity **(A)** and bar diagrams summarize quantification of the tumor flux with single datapoint for each mouse including mean and standard deviation of the group **(B)**. Kinetic of the tumor flux over time is shown for each mouse represented by one line **(C)**. n = 2 (naive), 4 (T cell), 4 (stCAR T cell).

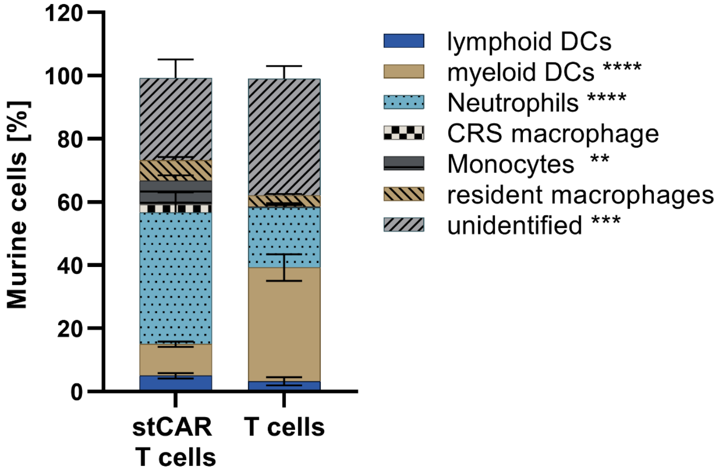


**Figure S3: Murine immune cell phenotyping in spleen.**

Tumor engrafted NSG-SGM3 mice treated with short-term (st) CAR T cell or T cell were sacrificed on the respective termination day and murine immune cells in the spleen were assessed by flow cytometry. Bar diagrams summarize the frequency of the different murine immune cells including mean and standard deviation. n = 4 (T cells), 4 (stCAR T cells). Statistics were determined by two-way ANOVA with Sidak's multiple comparisons test. **(p=0.0094), ***(p=0.0001), ****(p<0.0001).

### Supplemental Material and Methods

#### Murine immune cell phenotyping

Single cell suspension of splenocytes were stained for flow cytometry with anti-human CD45 (2D1, BV510, BioLegend) and the anti-mouse antibodies CD11b (M1/70, FITC, BioLegend), CD11c (N4/18, PE, eBioscience), F4/80 (BM8, BV605, BioLegend),Ly6C (1G7.G10, APC, Miltenyi Biotec), Ly-6G (REA526, PE-Cy7, Miltenyi Biotec) and viability was assessed with efluor780 (eFluor 780, Thermo Fisher Scientific). Mouse cells were determined as human CD45- and EBFP- cells. The mouse immune cells were determined as followed: Lymphoid DCs as CD11b-, CD11c+; myeloid DCs as CD11b+, CD11c+; neutrophils as CD11b+, CD11c-, Ly6G+; resident macrophages as CD11c-, F4/80+, Ly6C-; CRS-associated macrophages as CD11c-, F4/80+, Ly6C+; monocytes as CD11c-, F4/80-, Ly6C+; remaining cells were termed as unidentified.
